# Ancient and Animal-Specific Regulatory Modes of EWS::FLI1 Revealed by a Minimal Yeast Model

**DOI:** 10.1101/2025.10.22.680884

**Authors:** Diego Velázquez, Cristina Molnar, Jose Reina, Jaume Mora, Cayetano Gonzalez

## Abstract

Ewing sarcoma (EwS) is an aggressive, human-exclusive tumor typically driven by the EWS::FLI1 fusion protein. To assess whether the neomorphic functions of EWS::FLI1 are fundamentally dependent on evolutionarily recent cofactors such as ETS transcription factors (ETS-TFs), Plycomb group (PcG) proteins, CBP/p300, or specific subunits of the BAF complex, we expressed EWS::FLI1 in the model organism *Saccharomyces cerevisiae.* This minimal system was chosen because several key EWS::FLI1’s cofactors possess greatly reduced sequence homology (e.g., BAF) or are lacking altogether (e.g., ETS-TFs, PcG, or CBP/p300). We used co-IP/MS to map the yeast interactome, Chip-Seq to identify gDNA binding sequences, RNA-Seq for global gene expression, and engineered reporters to test conversion of (GGAA) tandem repeats (GGAAμSat) into neoenhancers. We found that the yeast EWS::FLI1 interactome was more limited and qualitatively distinct from its human counterpart, sharing core machinery (e.g. RNA Polymerase II, FACT) but lacking the BAF/SWI-SNF and spliceosome complexes, and showing strong enrichment for the SAGA chromatin remodeling complex. We also found that EWS::FLI1 binds to hundreds of sites in the yeast genome with a clear preference for putative ETS-TF consensus sequences and (CA) dinucleotide repeats. Yet, EWS::FLI1 expressing cells presented only minimal transcriptional dysregulation, a stark contrast to the extensive changes observed in humans and Drosophila cells. Finally, we found that EWS::FLI1 successfully converted silent GGAAμSat sequences into active enhancers in yeast. This remarkable result occurs despite the absence of homologs for key human activators, such as CBP/p300, strongly suggesting that EWS::FLI1 can mobilize functionally related, non-homologous pathways to establish neoenhancers at GGAAμSat sites. Altogether, our results indicate that EWS::FLI1’s core ability to drive GGAAμSat-dependent gene expression is a conserved, ancient property, while GGAAμSat-independent extensive transcriptome reprogramming is dependent on co-factors and pathways specific to animal cells.

## INTRODUCTION

Ewing sarcoma (EwS) is an aggressive, human exclusive tumor, typically arising in bone and soft tissues, that is reported to be the second most common bone malignancy in children, adolescents and young adults in the USA (Riggi *et al*, 2021). More than 80% of EwS patients express the EWS::FLI1 fusion protein that contains the transactivation domain of the FET family member EWS and the DNA binding domain of the E26 transformation specific domain transcription factor (ETS-TF) Friend leukemia integration 1 (FLI1). In one out of four EwS patients, the chimeric gene EWSR1-FLI1 encoding EWS::FLI1 is the only detectable genetic event at diagnosis (Crompton *et al*, 2014; Delattre *et al*, 1994; Sankar & Lessnick, 2011).

EWS::FLI1’s most conspicuous effect is massive reprogramming of gene expression that sets the stage for malignant transformation. Functional annotation of the EWS::FLI1 transcriptomic signature shows that upregulated genes are enriched for functions related to cell proliferation, energy metabolism and response to DNA damage, while downregulated genes are linked to differentiation and cell signaling (Cidre-Aranaz & Alonso, 2015; Grunewald *et al*, 2018; Kauer *et al*, 2009).

Well-documented EWS::FLI1’s direct target sequences include microsatellites made of GGAA repeats (GGAAμSats) and bona fide ETS-TF motifs which typically have a single or a small number of GGAA repeats (Gangwal *et al*, 2008; Guillon *et al*, 2009; Riggi *et al*, 2014). Upon binding to GGAAμSats, EWS::FLI1 recruits chromatin remodeling proteins, including the BAP complex, that convert these otherwise heterochromatic regions into euchromatic upstream activating sequences with enhancer or promoter capabilities (Boulay *et al*, 2017; Gangwal *et al*., 2008; Guillon *et al*., 2009; Patel *et al*, 2012; Riggi *et al*., 2014).

Given the vast number of sweet spot-sized GGAAμSats in the human genome—estimated to be in the order of tens of thousands (Johnson *et al*, 2017; Monument *et al*, 2014; Orth *et al*, 2022; Tomazou *et al*, 2015)—combined with the observation that nearly half of EWS::FLI1 binding sites are associated with these elements, and that GGAAμSats-driven ectopic enhancers can influence both proximal and distal genes (Johnson *et al*., 2017; Riggi *et al*., 2014), it is widely accepted that activation of GGAAμSats represents the primary mechanism by which EWS::FLI1 directly dysregulates gene expression (Gangwal *et al*, 2010; Gangwal *et al*., 2008; Guillon *et al*., 2009; Patel *et al*., 2012; Riggi *et al*., 2014). Direct experimental evidence demonstrating causation, i.e. inhibition of ectopic expression following targeted deletion of the respective microsatellite is still limited to few genes including *NR0B1*(*DAX1*) and *CAV1* (Gangwal *et al*., 2008; Guillon *et al*., 2009; Patel *et al*., 2012). However, an elegant experiment showing that zinc finger transcription factors engineered to repress transcription at GGAAμSats selectively silences a large fraction of the EwS gene expression program (Tak *et al*, 2022) substantiates the quantitative relevance of GGAAμSat-driven ectopic transcription in EwS.

The effect on ETS-TFs family binding motifs is more complex because EWS::FLI1 may either displace or form heterodimers with ETS-TFs (Guillon *et al*., 2009; Kauer *et al*., 2009; Riggi *et al*., 2014). Moreover, ETS-TFs can act as transcriptional repressors, activators, or both (Sharrocks, 2001).

In Drosophila, human EWS::FLI1 can convert transgenic GGAAμSat sequences into neoenhancers (Mahnoor *et al*, 2024; Molnar *et al*, 2022), but the Drosophila genome lacks natural sweet spot-sized GGAAμSats (Dmel ref genome r6.62). Yet, in Drosophila as in human cells, EWS::FLI1 expression brings about a massive dysregulation of transcription that affects hundreds of genes (Mahnoor *et al*., 2024; Molnar *et al*., 2022). Moreover, about a third of the upregulated genes included in two published EwS human signatures (Hancock & Lessnick, 2008; Kauer *et al*., 2009) are significantly enriched in the top of the GSEA-ranked Drosophila dataset (Molnar *et al*., 2022). In addition, the Drosophila EWS::FLI1 transcriptomic signature is highly significantly enriched for genes regulated by all eight ETS-TFs of Drosophila. Thus, human EWS::FLI1 can reproduce in flies two key neomorphic oncogenic functions: interfering with Drosophila’s own ETS-TFs as well as productively interact with the fly homologues of the proteins required for GGAAμSat conversion in human cells. Such a massive rewiring of the transcriptome caused by EWS::FLI1 in a species that lacks GGAAμSats shows that GGAAμSat-independent mechanisms can on their own have a very significant contribution to EWS::FLI1-induced transcriptional dysregulation (Molnar *et al*., 2022). This main conclusion underpins how objective limitations inherent to relatively simple study models like Drosophila can be exploited as experimental conditions (e.g. lacking GGAAμSats) that cannot be recreated in mammalian cells.

Building on this rationale, we chose to investigate the effects of EWS::FLI1 in a far more distantly related organism: *Saccharomyces cerevisiae*. Like Drosophila, the yeast genome lacks GGAAμSats. Moreover, unlike Drosophila, yeast lacks several key components known to be required for EWS::FLI1 neomorphic functions, like ETS transcription factors, Polycomb group (PcG) proteins, and the Nucleosome Remodeling and histone Deacetylase (NuRD) complex (Sharrocks, 2001), (Zhao *et al*, 2020) which are exclusive to metazoans. Furthermore, despite sharing common ancestry and critical core subunits, the mammalian BAF complex and its yeast counterpart SWI/SNF (SWItch/Sucrose Non-Fermenting) present notable differences including the loss of some components, the acquisition of novel ones, and substantial changes in subunit primary structure (Khavari *et al*, 1993; Lessard *et al*, 2007; Muchardt & Yaniv, 1993; Staahl *et al*, 2013; Wang *et al*, 1998; Wang *et al*, 1996a; Wang *et al*, 1996b). These distinctions raise the questions whether human EWS::FLI1 can mount a major dysregulation of the transcriptome, and drive GGAAμSat conversion, in yeast as it does in humans, or whether such neomorphic functions require animal cell-specific components that are absent or not sufficiently conserved in yeast.

## RESULTS

### The yeast EWS::FLI1 interactome reveals both common and distinct features compared to human and Drosophila

To identify the network of yeast proteins that human EWS::FLI1 might interact with, we performed co-immunoprecipitation (co-IP) using an anti-EWS antibody, followed by mass spectrometry (MS). We used DVS050 cells, a derivative of the W303-1A strain (see Material and Methods) transformed with pYES2 plasmids carrying EWS::FLI1 under the control of the pGAL1-inducible promoter. DVS050 cells transfected with empty pYES2 plasmid were used as control. We analysed samples taken after 6 hours of exposure to galactose. This analysis identified 54 yeast proteins as significant interaction partners of human EWS::FLI1 (fold change > 1.5, adjusted *p*-value < 0.05; Table S1) a considerably smaller number than observed in human and Drosophila cells (Elzi *et al*, 2014; Molnar *et al*., 2022; Selvanathan *et al*, 2015).

Functional annotation of this limited set of interacting proteins revealed only three significantly enriched clusters (Figure 1A). The two most prominent clusters encompass chromatin organization and regulation of transcription by RNA polymerase II (RNA Pol II), closely followed by RNA Pol II-dependent transcription (enrichment score 6.47 and 6.36, respectively). A third cluster, exhibiting a substantially lower enrichment score (1.74), is associated with TORC1 signalling.

**Figure 1.**
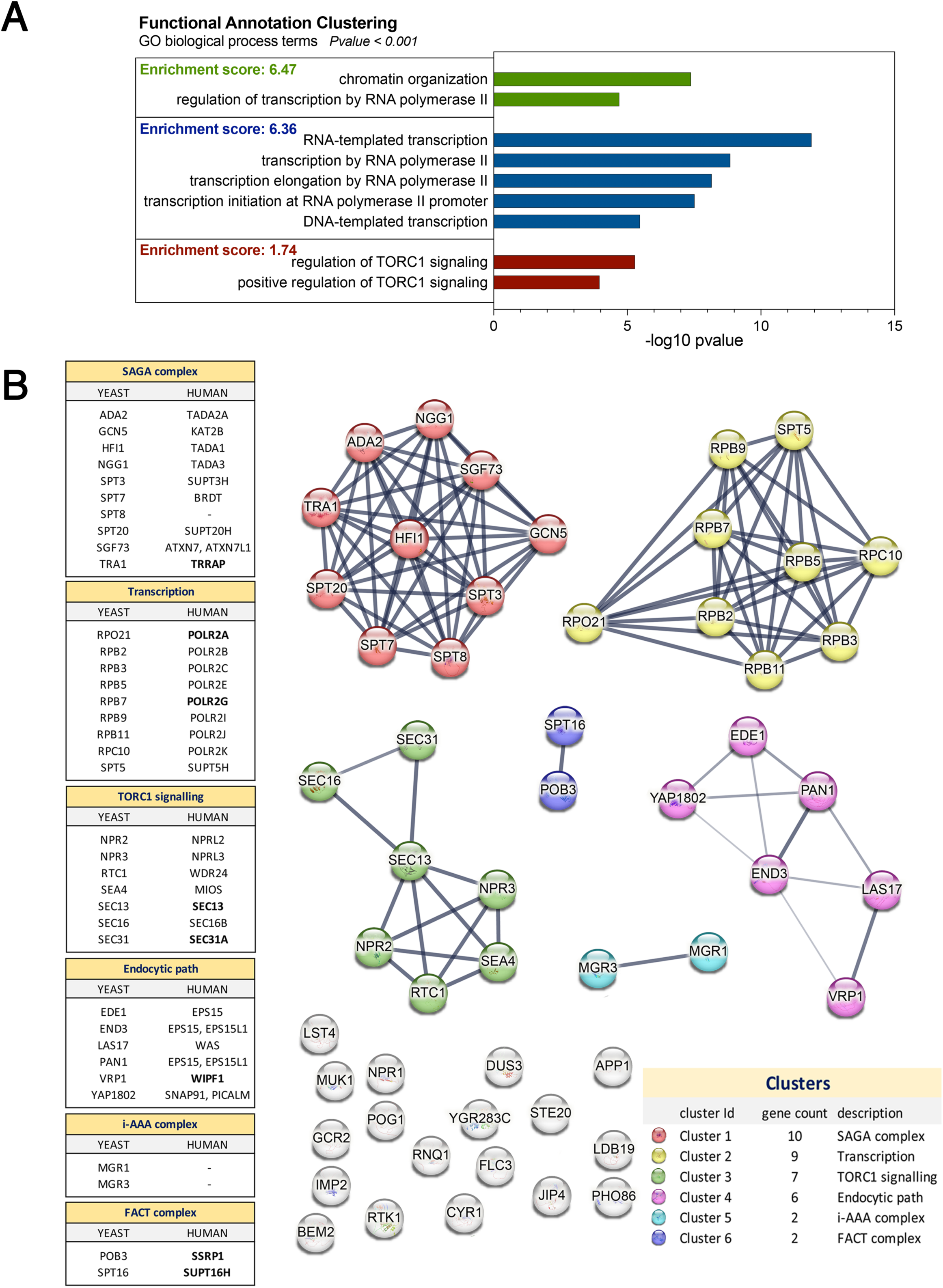
EWS::FLI1 interacts with protein complexes in yeast. **(A)** Functional annotation clustering of Gene Ontology (GO) biological process terms for the yeast proteins identified in the EWS::FLI1 interactome. Bar graphs represent the statistical significance of each GO term (−log10 *p*-value). **(B)** Network and cluster analysis of physical protein associations among EWS::FLI1 yeast interactors using the STRING database. Line thickness reflects the strength of data support. The tables on the left list the EWS::FLI1 yeast interactors and their human orthologs, grouped according to protein complexes. Human proteins in bold were found as EWS::FLI1 interactors in human cells (Elzi *et al*., 2014; Selvanathan *et al*., 2015). Data shown represent the mean of three independent biological replicates.

Network and clustering analysis using STRING showed that over half of the 54 EWS::FLI1-interacting proteins are organized into just six distinct networks of verified physical protein-protein interactions (Figure 1B). These include: 10 components of the SAGA (Spt–Ada–Gcn5–acetyltransferase) complex; 9 subunits of RNA polymerase, comprising the RNA Pol II catalytic core and subunits shared across RNA polymerases I, II, and III; 5 components of the Seh1-associated (SEA) complex, along with 2 additional proteins involved in Coat Protein Complex II (COPII) vesicle formation at ER exit sites; and 6 proteins implicated in clathrin-mediated endocytosis and the coordination of actin polymerization with vesicle formation. Additionally, four other EWS::FLI1 interactors form two smaller, two-component networks: one consisting of the nuclear-encoded components of the mitochondrial inner membrane i-AAA Protease complex, Mgr1 and Mgr2, and the other comprising the two-component *Saccharomyces cerevisiae* FACT (Facilitates Chromatin Transcription) complex, Pob3 and Spt16.

Broadly, these STRING networks fall into two major functional groups. The first is linked to transcription and includes SAGA, RNA Pol II, and FACT. The SAGA complex acts as a transcriptional co-activator, primarily through chromatin modification and interaction with the basal transcription machinery, and is crucial for the expression of numerous yeast genes, particularly those involved in stress response. RNA Pol II is responsible for transcribing protein-coding genes as well as most small nuclear RNAs (snRNAs) and microRNAs (miRNAs), playing a central role in yeast cellular processes and adaptation. The FACT complex is a histone chaperone that facilitates the dynamic reorganization of nucleosomes necessary for efficient RNA Pol II-dependent transcription.

The second group is associated with vesicle trafficking and nutrient sensing, encompassing the STRING networks related to COPII, SEA, and clathrin-mediated endocytosis. The COPII coat complex forms vesicles at ER exit sites (ERES) and is essential for the transport of newly synthesized proteins from the endoplasmic reticulum (ER) to the Golgi apparatus, a critical step in the secretory pathway. The COPII complex is connected to SEA through the shared Sec13 protein. SEA acts as a GAP (GTPase-activating protein) for the Rag GTPase Gtr1, which plays a vital role in nutrient sensing and regulating the Target of Rapamycin Complex 1 (TORC1) signaling pathway. Notably, SEA and TORC1 physically interact and co-localize to the vacuolar membrane (the yeast equivalent of the mammalian lysosome). Clathrin-mediated endocytosis (CME) is primarily involved in internalizing cargo from the plasma membrane into the cell and in transport from the trans-Golgi network (TGN) to endosomes. It utilizes clathrin protein to form a coat around vesicles. Indeed, the endpoint of CME is the vacuole, where mature endosomes deliver their contents for degradation and recycling. Importantly, the vacuole’s functional state influences the dynamics and efficiency of CME, a process that appears to involve the SEA complex and TORC1 signaling pathway. Of note, nearly half of the remaining EWS::FLI1 interactors, which are not clustered by STRING, are nonetheless known to function in TORC1 signaling, stress/nutrient sensing and adaptation (Npr1, Lst4, Cyr1, Pog1), endosomal and secretory trafficking, and vacuolar protein sorting (Jip4, Ldb19 (Art1), Muk1, App1, Pho86) (Figure 1B).

These results show that human EWS::FLI1 can interact with certain protein complexes in yeast, including, notably histone chaperones required for RNA Pol II as well as RNA Pol II itself that are known interactors of EWS::FLI1 in human EwS cells as well as in Drosophila. However, the yeast EWS::FLI1 interactome is much smaller and presents notable qualitative differences with both the human and Drosophila interactomes, such as the absence of spliceosome and SWI/SNF proteins as well as, prominently, the enrichment of SAGA complex proteins.

### *EWS::FLI1* expression induces minimal transcriptional dysregulation

To investigate the putative effect of the human EWS::FLI1 oncogene on gene expression in yeast, we carried out RNA-seq on DVS050 cells transformed with pYES2 plasmids carrying EWS::FLI1. DVS050 cells transformed with either empty pYES2 plasmid or pYES2 plasmids carrying the DNA-binding mutant variant EWS::FLI1_RRLL_ were used as control. EWS::FLI_RRLL_ is a double mutant version (R361L+R364L) of EWS::FLI1 that prevents DNA binding (Bailly *et al*, 1994). Expression of this variant in *S. cerevisiae* shows no toxicity, with growth levels similar to those of yeast transformed with the empty pYES2 plasmid (Supplemental Figure 1).

We reasoned that with a cell cycle length of about 2 hours under these conditions, a 3-hour time point should suffice to observe most of the changes in the transcriptome caused by EWS::FLI1 expression. In addition, to get a fuller picture of the putative EWS::FLI1 effects, we also analysed samples taken after 6 hours (i.e about 3 consecutive cell cycles) of exposure to the oncogene).

Principal component analysis of the entire dataset shows that cells expressing EWS::FLI1 for 6 hours cluster apart while differences among all others experimental conditions are minor (Figure 2A). Indeed, the effect of EWS::FLI1 at 3 hours after induction is limited to 23 upregulated and 4 downregulated transcripts compared to control cells (Figure 2B,C; Table S2). At this point, the effect of the DNA binding mutant EWS::FLI1_RRLL_ is negligible with only 5 transcripts upregulated and none significantly downregulated (Figure 2B,C; Table S2). The very limited effect of EWS::FLI1 upon 3 hours of expression strongly suggests that the number of direct targets of this oncogene in yeast is very small.

**Figure 2.**
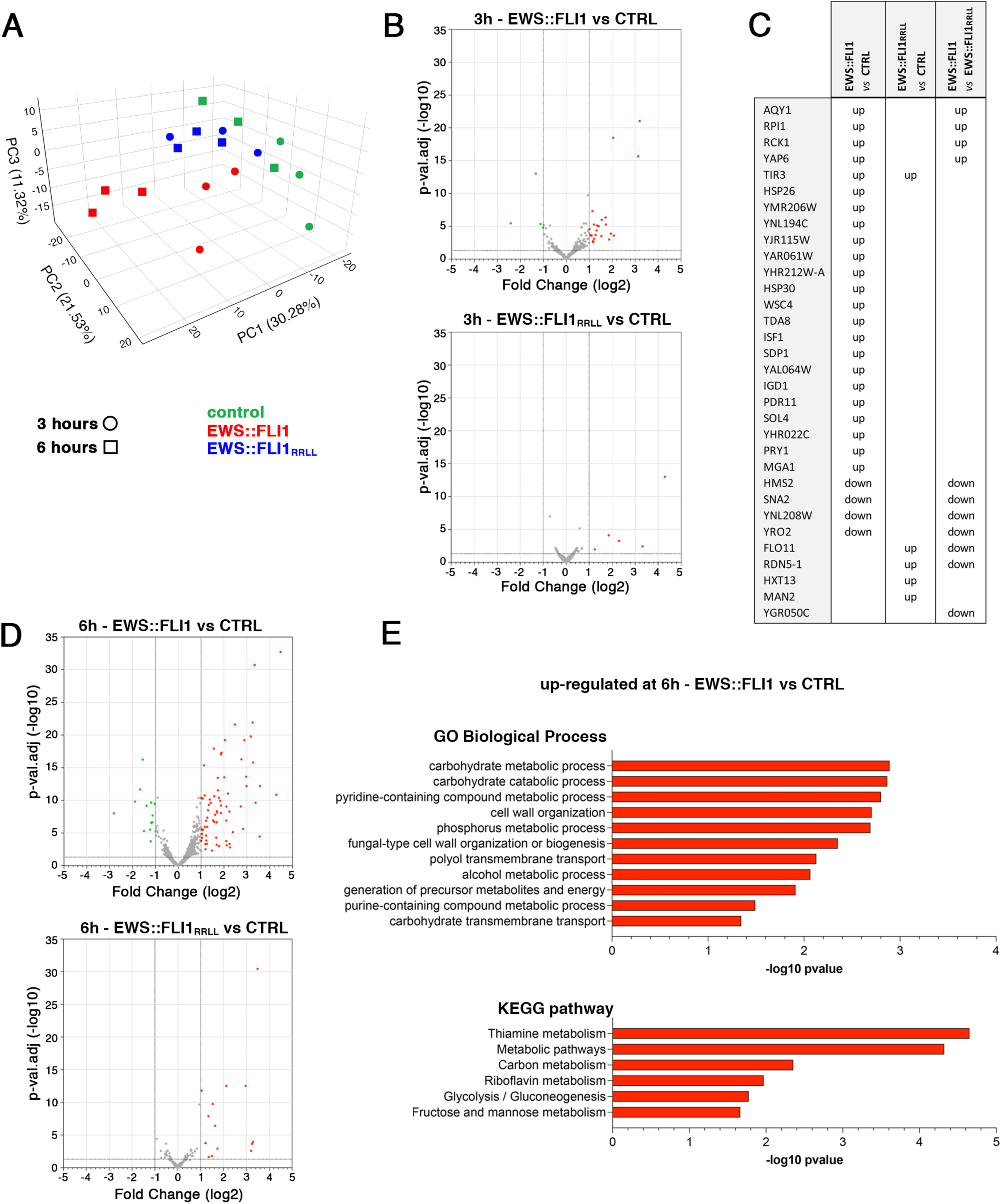
EWS::FLI1 expression dysregulates the yeast transcriptome. **(A)** Principal component analysis (PCA) RNA-seq data from cells expressing EWS::FLI1 (orange) or EWS::FLI1_RRLL_ (blue) and control cells (green), at 3 hours (circles) and 6 hours (squares) of induction. **(B)** Volcano plots showing changes in gene expression levels in EWS::FLI1, and EWS::FLI1_RRLL_ expressing cells compared to control samples, at 3 hours of induction. Red and green dots represent genes that are significantly (*p*-val.adj < 0.05) up (FC > 2) or downregulated (FC < -2) genes, respectively. **(C)** List of dysregulated genes in EWS::FLI1 and/or EWS::FLI1_RRLL_ expressing cells at 3 hours of induction. **(D)** Volcano plots showing changes in gene expression levels in EWS::FLI1, and EWS::FLI1_RRLL_ expressing cells compared to control samples, at 6 hours of induction. Red and green dots represent genes that are significantly (*p*-val.adj < 0.05) up (FC > 2) or downregulated (FC < -2) genes, respectively. **(E)** Bar graphs showing significantly (*p*-value < 0.05) enriched GO Biological Process terms and KEGG pathways of upregulated genes at 6 hours of EWS::FLI1 induction. All results shown in this figure were obtained from three independent biological replicates.

Transcriptome changes at 6 hours are more abundant, but still relatively minor with only 73 and 13 genes significantly up and downregulated, respectively (Figure 2D; Table S2). Among them are 20 transcripts that were upregulated and 3 of the transcripts that were downregulated at the 3-hour time point. Consistently with its lack of effect on yeast viability, the effect of EWS::FLI1_RRLL_ is also negligible at the 6-hour time point with only 18 upregulated transcripts and none significantly downregulated. Interestingly, the overlap in transcriptional responses to EWS::FLI1 and EWS::FLI1_RRLL_ is minimal: only 1 and 12 transcripts upregulated by both variants at 3 and 6 hours, respectively. These results suggest that the effect of EWS::FLI1_RRLL_ is not only quantitatively smaller, but also qualitatively different than that of EWS::FLI1.

Functional annotation using the DAVID web tool of the 6h EWS::FLI1 yeast transcriptome identified no significantly (*p*-value < 0.05) enriched Gene Ontology (GO) terms and KEGG pathways among downregulated genes and only a small number of among upregulated genes. This was to be expected given the very low number of genes involved. Most of the enriched terms and pathways were related to metabolic processes and cell wall organisation and a substantial fraction of the affected transcripts overlaps with published transcriptomics data on *S. cerevisiae*’s response to nutritional changes, ionic stress, salt stress, oxidative stress, osmotic shock, and genetic interventions including gene deletion and overexpression (Figure 2E) (Gasch, 2003; Taymaz-Nikerel *et al*, 2016). These results suggest that most of the genes dysregulated at the 6-hour time point reflect the yeast cells response to EWS::FLI’s toxic effects.

In summary, our results show that the number of dysregulated transcripts observed even after 6 hours of EWS::FLI1 expression in yeast is an order of magnitude smaller than the established EWS::FLI1 signatures in human signatures in human (Hancock & Lessnick, 2008; Kauer et al., 2009) and Drosophila (Mahnoor et al., 2024; Molnar et al., 2022). From these observations, we conclude that in stark contrast to the extensive transcriptional changes observed in animal cells of different species, *EWS::FLI1* expression induces minimal transcriptional dysregulation in yeast.

### EWS::FLI1 binds to multiple chromatin sites causing a major relocalisation of RNA polymerase II in *S. cerevisiae*

To determine if human EWS::FLI1 binds to yeast chromatin, we used chromatin immunoprecipitation followed by DNA sequencing (ChIP-seq) with an anti-EWSR1 antibody. In parallel, we profiled genome-wide RNA Pol II occupancy using an anti-RNA Pol II antibody, while an anti-GFP antibody served as negative control. All experiments were conducted in DSVS050 cells transformed with either a plasmid expressing EWS::FLI1 or the empty plasmid, under the same RNA-seq induction conditions.

We identified 834 high-confidence EWS::FLI1 peaks in EWS::FLI1-expressing yeast cells, mapping to 788 annotated genes. Genomic annotation revealed a strong preference for gene-associated regions, with nearly half of the peaks (47.1%) found at promoters, and most of the remaining peaks distributed between transcription termination sites (TTS) and exons (21.2% and 27.2%, respectively) (Figure 3A). Only 4.4% of peaks were in intergenic regions, which are therefore markedly underrepresented since they account for about 27% of the *S. cerevisiae* genome.

**Figure 3.**
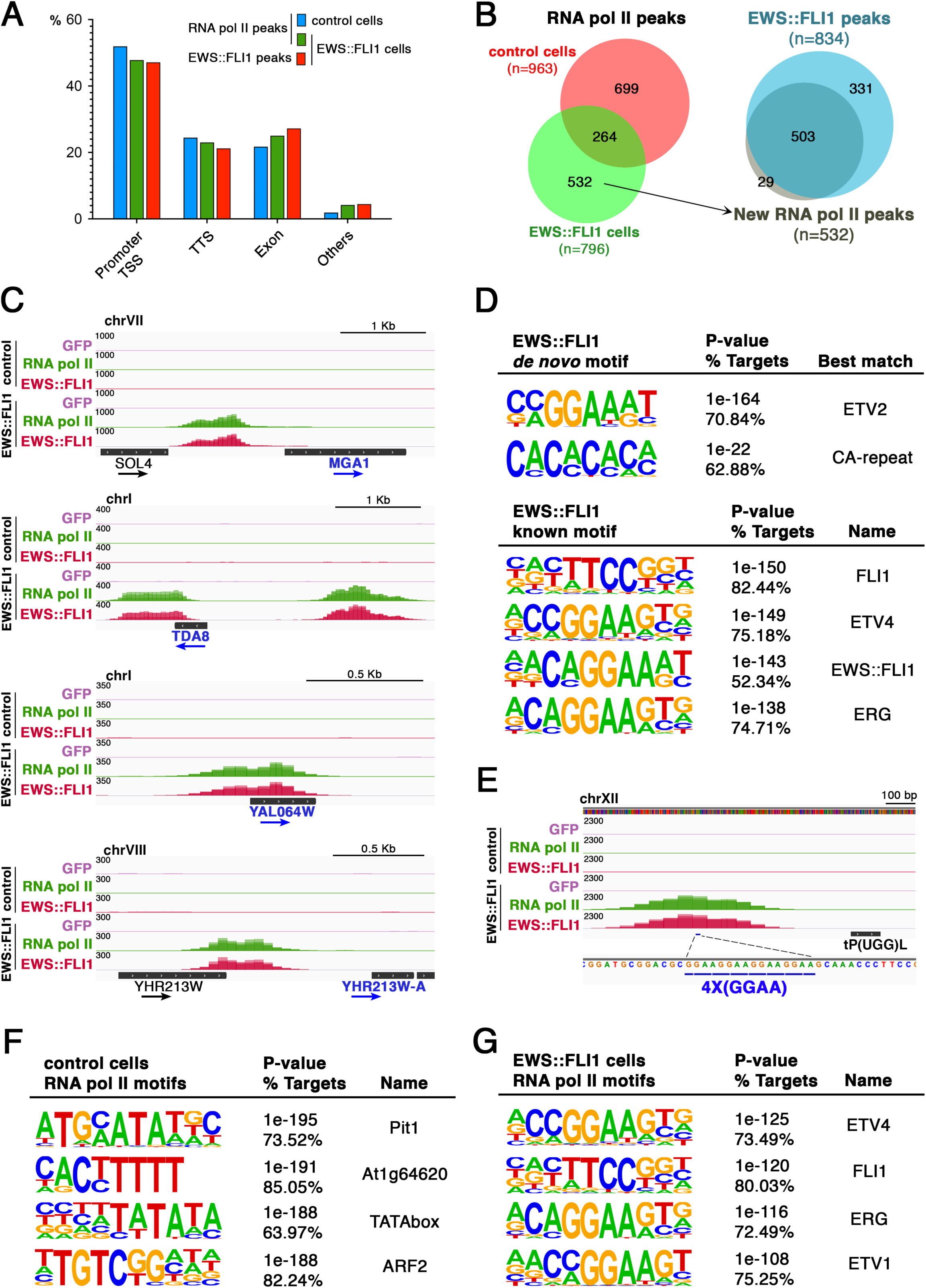
EWS::FLI1 binds chromatin and redistributes RNA polymerase II in *S. cerevisiae*. **(A)** Genomic distribution of RNA Pol II peaks (blue and green) and EWS::FLI1 peaks (red) in control (blue) or in EWS::FLI1-expressing cells (green and red). Peaks were annotated relative to promoters and transcription start sites (TSS), transcription termination sites (TTS), exons, and other regions (introns and intergenic regions). **(B)** Venn diagrams showing the overlap between RNA Pol II peaks in control cells (n = 963) and in EWS::FLI1-expressing cells (n = 796), together with EWS::FLI1 peaks (n = 834). Of the 532 newly acquired RNA Pol II peaks in EWS::FLI1-expressing cells, 95% overlapped with EWS::FLI1 binding sites. **(C)** Genome browser tracks showing EWS::FLI1 and RNA Pol II occupancy in control and EWS::FLI1-expressing cells at the loci corresponding to the four upregulated genes (gene names in blue). RNA Pol II (green), EWS::FLI1 (red), and negative control GFP (grey) ChIP-seq signals are shown. **(D)** Motif analyses of EWS::FLI1 binding sites. *De novo* motif analysis identified a consensus ETS motif (CCGGAAAT, top) and CA-dinucleotide repeats (bottom). Known motif analysis confirmed the enrichment of ETS-family motifs (top four motifs shown). **(E)** Genome browser view showing an EWS::FLI1 and RNA Pol II peak at the only 4×(GGAA) repeat present in the *S. cerevisiae* reference genome. RNA Pol II (green), EWS::FLI1 (red), and GFP (grey) ChIP-seq signals are shown for control and EWS::FLI1-expressing cells. **(F)** Motif enrichment at RNA Pol II peaks in control cells. The top four motifs are shown. **(G)** Motif enrichment at RNA Pol II peaks in EWS::FLI1-expressing cells. The top-ranked motifs corresponded to ETS-family transcription factors (ETV4, FLI1, ERG, ETV1), consistent with relocalization of RNA Pol II to EWS::FLI1-bound loci. All results shown are based on three independent biological replicates.

Analysis of RNA Pol II binding revealed 963 peaks in control cells and 796 peaks in EWS::FLI1-expressing cells, with 264 sites shared between conditions (Figure 3B). Notably, 75% of the EWS::FLI1 peaks (626/834) overlapped with RNA Pol II peaks in EWS::FLI1-expressing cells. This overlap corresponded with a pronounced redistribution of RNA Pol II occupancy: of the 963 sites detected in control cells, 699 were lost, while 532 new sites emerged, upon EWS::FLI1 expression. Strikingly, 95% of these newly acquired RNA Pol II peaks overlapped with EWS::FLI1 binding sites, indicating that polymerase recruitment largely coincides with EWS::FLI1 target loci (Figure 3B). In terms of genomic distribution, RNA Pol II peaks in control cells were localized as 52% at promoters, 24.5% at TTS, 21.7% within exons, 0.3% within introns, and 1.6% in intergenic regions. In EWS::FLI1-expressing cells, RNA Pol II peaks were distributed as 47.8% at promoters, 23% at TTS, 25% within exons, and 4.2% in intergenic regions (Figure 3A). These results indicate that both EWS::FLI1 and RNA Pol II preferentially occupy promoter-proximal regions, and that RNA Pol II redistribution largely reflects the genomic localization of EWS::FLI1. Together, these findings demonstrate that EWS::FLI1 binds directly to hundreds of chromatin sites in *S. cerevisiae* and induces a broad redistribution of RNA Pol II, with polymerase occupancy largely redirected to EWS::FLI1 binding sites.

The observed RNA Pol II relocalization is somewhat unexpected, as it is not accompanied by widespread transcriptional dysregulation. Indeed, most RNA Pol II sites lost (699/963) and gained (532/796) upon EWS::FLI1 expression did not correspond to genes dysregulated at either the 3- or 6-hour time points. Furthermore, only a minority of the genes upregulated (4/23 at 3 hours and 17/73 at 6 hours) and none of the genes downregulated harbored EWS::FLI1 binding sites. Notably, the four genes upregulated at 3 hours exhibited the strongest EWS::FLI1 peaks that also coincided with strong RNA Pol II peaks, suggesting that these may represent direct transcriptional targets of EWS::FLI1 in *S. cerevisiae* (Figure 3C).

To further characterize EWS::FLI1 binding sites, we performed motif analyses using HOMER. *De novo motif analysis* identified a consensus ETS binding sequence (5′-CC<u>GGAA</u>AT-3′) as the most significantly enriched motif at EWS::FLI1 peaks (present in 71% of bound regions compared to 26% of background sequences; *p*-value = 1e−164). Additionally*, known motif analysis* further confirmed the strong enrichment of ETS-family motifs (Figure 3D) like those associated with human FLI1, ETV4, ERG, ELK4, ETV1, and ETS1 (present in more than 60–80% of target sequences compared to only 20–35% in the background; *p*-value < 1e-120). The EWS::FLI1 fusion motif itself (xACAGGAAAT) was also highly enriched (*p*-value = 1e−143, 52% vs. 15%). Interestingly, the four genes upregulated at 3 hours that exhibited the strongest EWS::FLI1 peaks referred to earlier also harbour core GGAA motifs.

A second enriched motif (present in 63% of bound regions versus 46% in background sequences; *p*-value = 1e−22), corresponded to 10x(CA) repeats. In addition, we also detected enrichment of regulatory sequences bound by other transcription factors such as E2F4 (28% vs. 10%; *p*-value = 1e−49) and TFDP1 (20% vs. 6%; *p*-value = 1e−37), as well as composite ETS:IRF motifs like PU.1–IRF (50% vs. 29%; *p*-value = 1e−35). A canonical TATA-box motif (41% vs. 25%; *p*-value = 1e−24) was also enriched, consistent with the promoter-proximal localization of many EWS::FLI1 peaks (Supplemental Figure 2).

GGAA repeats are extremely rare in the *S. cerevisiae* reference genome: only four 3×(GGAA) and a single 4×(GGAA). We did not detect EWS::FLI1 peaks at any of the 3×(GGAA) repeats, but a clear EWS::FLI1 peak was observed at the 4×(GGAA) repeat (Figure 3E).

*De novo* and known motif analysis of RNA Pol II peaks identified TATA-box motifs and other transcription factor consensus sequences (Figure 3F). Notably, in the presence of EWS::FLI1, the top-ranked RNA Pol II-associated motifs corresponded to ETS-binding sites, highlighting that EWS::FLI1 relocalised RNA Pol II to EWS::FLI1-bound regions, predominantly containing putative ETS binding motifs (Figure 3G).

Collectively, our findings demonstrate that EWS::FLI1 preferentially targets multiple promoter-proximal chromatin sites in S*. cerevisiae*, with a distinct preference for sequences containing putative ETS binding motifs. The binding of EWS::FLI1 at these loci leads to the redirection of RNA Pol II occupancy; however, this relocalization results in the transcriptional activation of only a subset of these target genes.

### EWS::FLI1 reverts GGAAμSat-dependent chromatin silencing in yeast

In sharp contrast to humans, there are no GGAAμSats in *S. cerevisiae* (R64-5-1 reference genome). Therefore, the first step to test the effect of EWS::FLI1 on these satellite sequences in yeast was to create new strains carrying engineered reporters of GGAAμSat-driven transcription like those that have been used in Drosophila and human cells (Holting *et al*, 2022; Molnar *et al*., 2022). These reporter constructs carry a gene that encodes a fluorescent protein under the control of a modified constitutive promoter, and is inserted downstream of a tandem array of GGAA repeats. Under these conditions, transcription of the fluorescent reporter is normally downregulated by the proximity to the GGAA repeats, but becomes upregulated upon expression of EWS::FLI1.

Following this approach, we used a modified version of the iso-1-Cytochrome C constitutive promoter (*CYC1p*) (Zekhnini *et al*, 2023), which is widely used as a template for synthetic promoters (Guarente *et al*, 1984; Guarente *et al*, 1982; Hahn *et al*, 1985). Reporter strains were made by integrating constructs carrying 10x(GGAA), 20x(GGAA), or 40x(GGAA) repeats upstream the *CYC1p*-GFP sequence from the pRS426 plasmid (Zekhnini *et al*., 2023) into the *TRP1* gene of W303-1A wild-type cells (Figure 4A). Similar strains carrying a random 80-nucleotide insertion or no insertion at all upstream of the *CYC1p*-GFP sequence were also built as control cells (Figure 4A; Supplemental Figure 3).

**Figure 4:**
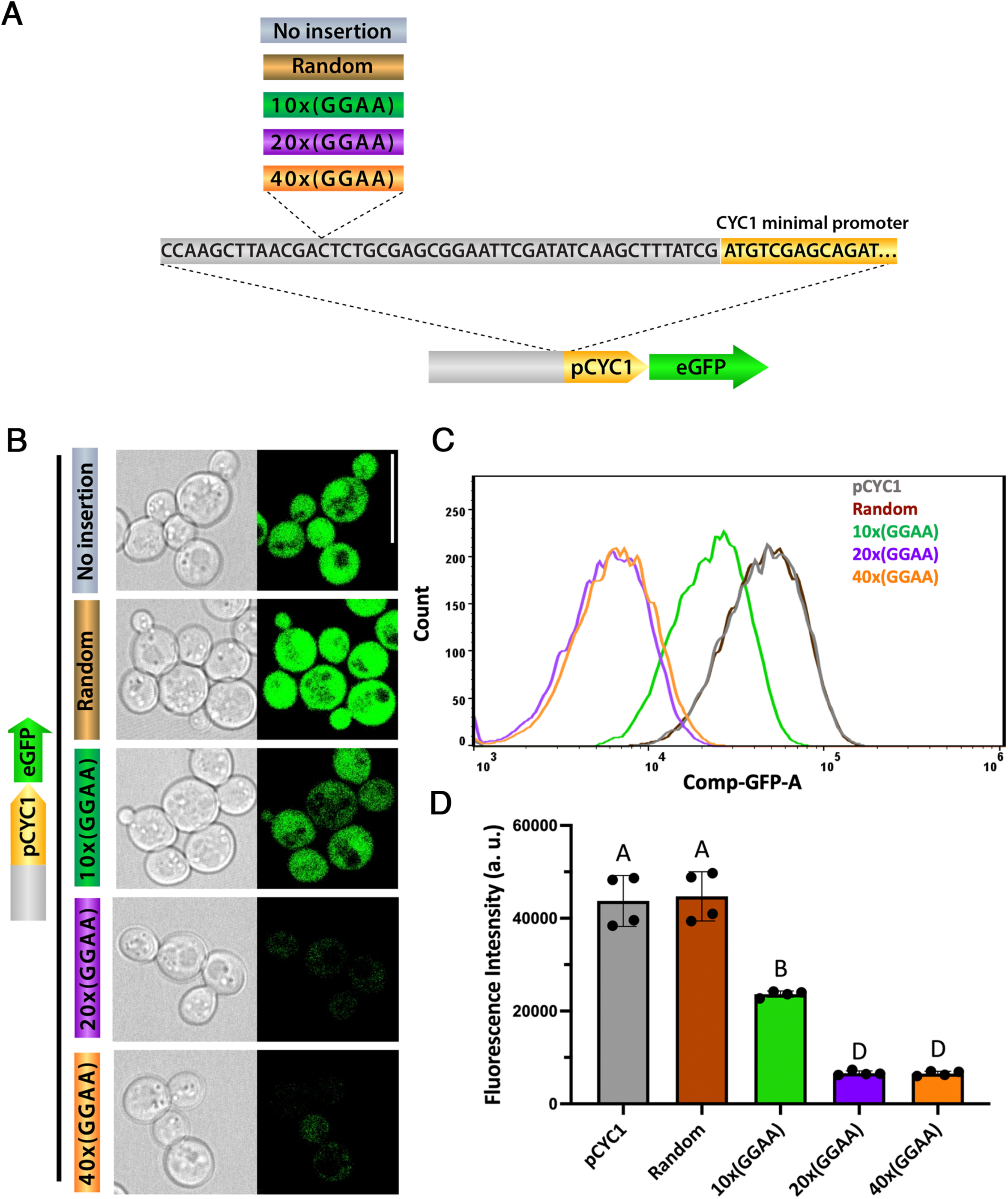
Introduction of human GGAA microsatellites into the yeast genome induces gene silencing. **(A)** Schematic depicting of the constructs carrying eGFP driven by the minimal *pCYC1* promoter with upstream inserts of an 80-nt random or 10×/20×/40×(GGAA) repeats. **(B)** Confocal microscopy images of W303-1A isogenic derivatives containing *pCYC1* with 80-nt random or 10×/20×/40×(GGAA) inserts. Cells were grown in Ura^−^Raff to OD_600_ = 0.2 and induced with 2% galactose for 6 h. Representative GFP fluorescence (right) and bright-field images showing whole-cell morphology (left) are shown. Scale bar, 10 μm. **(C-D)** Quantification of GFP expression by FACS. Cells (n= 10,000) cells were cultured under the same conditions as in (B). (C) Histogram showing the distribution of cell counts as a function of Comp-GFP. (D) Quantification of eGFP fluorescence. Mean +/-SD from three independent replicates are represented. Letters indicate statistically significant differences between means (one-way ANOVA, *p*-value < 0.05). a.u., arbitrary units.

We observed nearly equally strong eGFP expression both in cells transformed with the *CYC1p*-GFP plasmid that carries no GGAA repeats (4.35E+04 ±5.48E+03) and in cells transformed with the *CYC1p*-GFP plasmid carrying a random, 80-nucleotide insertion minor (4.45E+04 ±5.30E+03) (Figure 4B-D, grey and brown, respectively). Difference in eGFP level between these two conditions was not significant (*p*-value = 0.9995) thus showing that the random 80-nucleotide insertion has no notable effect on eGFP expression (Figure 4B-D). In contrast, eGFP expression is reduced in cells carrying the 10x(GGAA)μSatellite (2.37E+04 ±6.57E+02; *p*-value < 0.0001; Figure 4B-D, green), and much more so in cells carrying the 20x and 40x(GGAA)μSatellites, (6.60E+03 ±4.87E+02 and 6.54E+03 ± 4.94E+02, respectively; Figure 4B-D, purple and orange, respectively). Differences between the 10x(GGAA) and both the 20x(GGAA) and 40x(GGAA) insertions are also highly significant (*p*-value < 0.0001) while difference between 20x(GGAA) and the 40x(GGAA) are not (*p*-value = 0.9998) (Figure 4B-D).

These results are consistent with published data showing that in *S. cerevisiae*, as in mammalians, an expanded array of a low complexity repetitive DNA sequence is sufficient to direct the assembly of a silent heterochromatic chromosomal domain (Loo & Rine, 1994; Stavenhagen & Zakian, 1994). They also show that 20x(GGAA)μSats and 40x(GGAA)μSats are much more effective than 10x(GGAA)μSats as far transcriptional repression of nearby genes is concerned, while there are no significant differences in efficiency between 20x and 40x(GGAA)μSats.

We then tested the ability of the human EWS::FLI1 oncogene to bring about transcription from the *CYC1p* promoter silenced by proximity to (GGAA)μSats. To this end, we transformed our W303-1A derivative strains carrying *CYC1p*-GFP, 20x(GGAA)μSat-*CYC1p*-GFP and 40x(GGAA)μSat-CYC1p-GFP with pYES2 plasmids carrying EWS::FLI1. Similar transformants created with pYES2 plasmids that were either empty or expressing the DNA-binding mutant variant, EWS::FLI1_RRLL_, were used as controls (Supplemental Figure 4).

We found that 6 hours after exposure to galactose, eGFP fluorescence levels were indistinguishable high in CYC1p-GFP cells transformed with pYES2 plasmids that were empty (5.32E+04 ±1.44E+03) or expressing either EWS::FLI1_RRLL_ (5.17E+04 ±5.12E+03) or EWS::FLI1 (5.27E+04 ±2.21E+03) (Figure 5A, upper panel). As expected, eGFP fluorescence was strongly downregulated in 20x(GGAA)μSat-CYC1p-GFP stable transformants transfected with either empty pYES2 plasmids (2.82E+03 ±7.30E+02; *p*-value < 0.001) and so it was upon transfection with pYES2 plasmids expressing EWS::FLI1_RRLL_ (4.80E+01 ±1.33E+03; *p*-value < 0.001) (Figure 5A, middle panel). Conversely, eGFP fluorescence was significantly recovered in 20x(GGAA)μSat-CYC1p-GFP cells expressing EWS::FLI1 via pYES2 transformation (1.40E+04 ±1.10E+03; *p*-value < 0.001) (Figure 5A, middle panel). Similar results were observed in 40x(GGAA)μSat- CYC1p-GFP stable transformants (Figure 5A, bottom panel). Notably, this recovery appears strictly dependent on the oncogene’s prion-like domain because EWS::FLI1_YS37,_ a mutant variant that replaces all 37 tyrosine residues essential for condensate assembly in human cells (Boulay *et al*., 2017) failed to rescue eGFP fluorescence in 20x(GGAA)μSat-CYC1p-GFP cells (Supplemental Figure 1).

**Figure 5:**
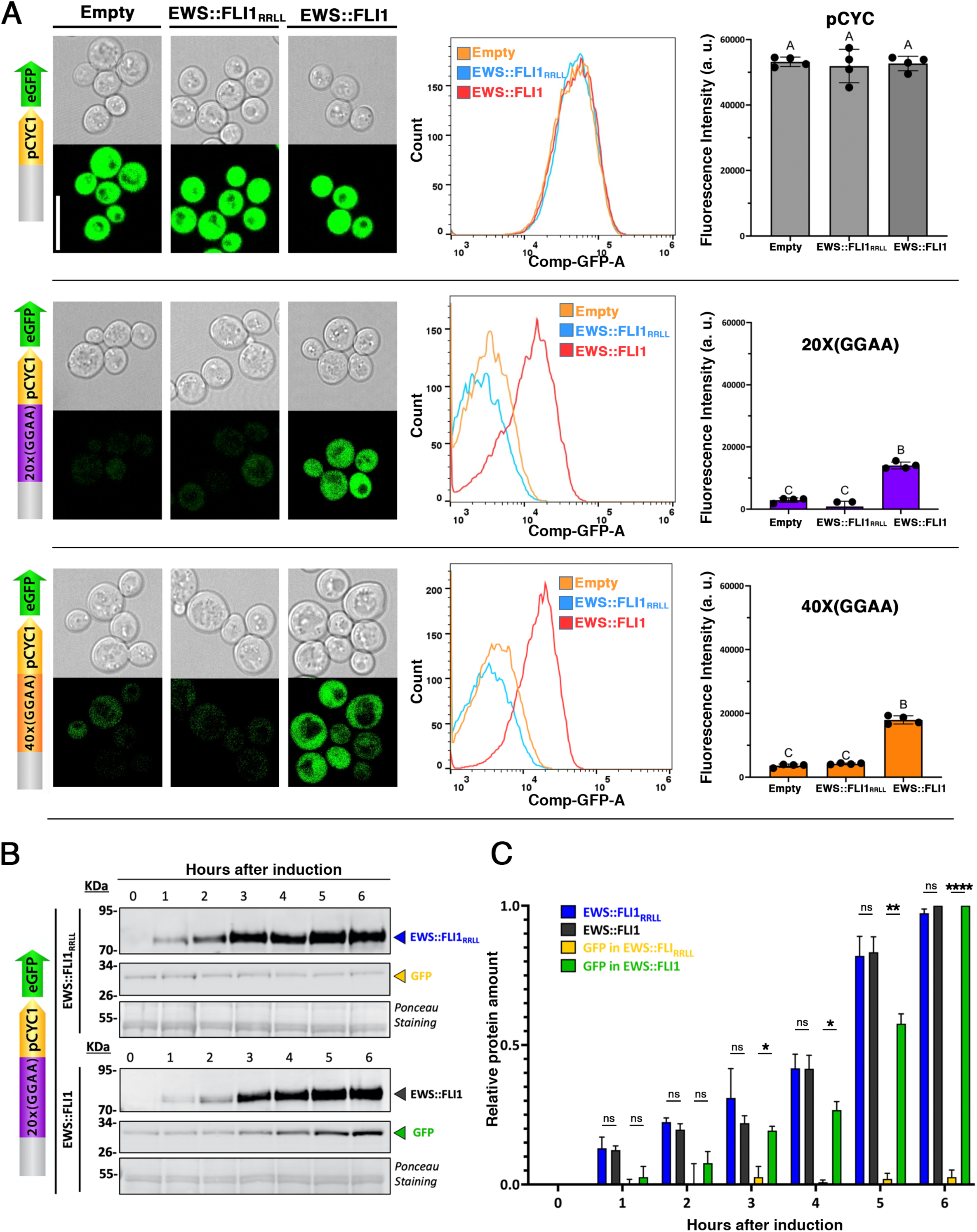
Transcriptional activation by EWS::FLI1 through recognition of GGAA microsatellites in yeast. **(A)** W303-1A derivatives carrying pCYC1 (top), 20×(GGAA) (middle), or 40×(GGAA) (bottom) reporters were transformed with empty pYES2, EWS::FLI1, or EWS::FLI1_RRLL_. Cells were cultured in Ura^−^Raff medium to OD_600_ = 0.2, induced with 2% galactose for 6 h, and harvested. Representative confocal images (left panels) showing bright-field (top) and GFP fluorescence (bottom) are shown. FACS analysis of GFP fluorescence is shown as histograms (middle panels) representing cell count distributions as a function of Comp-GFP, and as bar charts (right panels) showing the mean +/-SD of three independent replicates. Letters indicate statistically significant differences between means (one-way ANOVA, *p*-value < 0.05). a.u., arbitrary units. **(B)** Time-course analysis of 20x(GGAA)_*pCYC1*_eGFP cells transformed with pYES2 EWS::FLI1_RRLL_ (top) or EWS::FLI1 (bottom). Cells were grown in YNB lacking uracil supplemented with 2% raffinose (Ura^-^ Raff) until OD_600_= 0.25 and then 2% of galactose was added. Samples collected at the indicated times were analyzed by Western blot with anti-EWSR1 and anti-GFP monoclonal antibodies to detect EWS::FLI1 and GFP, respectively. Arrowheads indicate EWS::FLI1 and GFP signal. Ponceau staining is shown as a loading and transfer control. The same results were obtained in two other Western blots carried out using independent biological replicates. **(C)** Relative quantification of EWS::FLI1 and GFP expression levels, normalized to 100% based on the mean signal of three independent experiments for EWS::FLI1 expression at 6 hours of induction. Error bars indicate SD. *p<0.05; ns = no significant; two-way ANOVA. a.u., arbitrary units.

A time-course analysis of 20x(GGAA)μSat-CYC1p-GFP W303-1A (DVS050) cells by Western blot shows the concomitant buildup of both EWS::FLI1 and eGFP over the 6-hour period upon the induction of EWS::FLI1 expression with galactose (Figure 5B,C) and confirms that eGFP fluorescence remains at background level despite increasing levels of the EWS::FLI1_RRLL_ DNA-binding mutant (Figure 5B,C). Importantly, the steady upregulation of GFP expression during the 6-hour period, which is already apparent at 3 hours, substantiates our reasoning that these time points should suffice to observe changes in the transcriptome caused by EWS::FLI1.

These results show that human EWS::FLI1 can productively interact with and convert transcription silencing GGAAμSats into neoenhancers.

## DISCUSION

To delineate the molecular requirements for the neomorphic transforming functions of the EWS::FLI1 oncoprotein, we have investigated its activity in a minimal eukaryotic system. Specifically, we sought to determine if these functions are fundamentally dependent upon evolutionarily recent cofactors, including ETS transcription factors, Polycomb Group (PcG) proteins, the p300 histone acetyltransferase, or the BAF chromatin remodeling complex. We have addressed this by expressing EWS::FLI1 in the model yeast, *Saccharomyces cerevisiae*. This organism lacks direct orthologs for several of these mammalian cofactors or possesses others with significantly divergent conservation, thus providing a “stripped-down” cellular environment to probe EWS::FLI1’s activities. In particular, we have investigated the effect of the human oncogenic fusion EWS::FLI1 in *Saccharomyces cerevisiae* with regards to proteomic interactions, DNA binding sites, its effect on the transcriptome, and its potential to drive gene expression from otherwise silent GGAAμSat sequences.

We have found that while the human and Drosophila EWS::FLI1 interactomes are strikingly similar (Molnar *et al*., 2022), the yeast interactome presents both notable commonalities but also substantial distinctions to both. The primary shared feature across all three species is the significant enrichment of RNA polymerase II (including the catalytic and other subunits) and its facilitator, FACT. In addition, terms associated with vesicle-mediated transport are also significantly enriched in the interactomes from all three species, although to a much higher level in yeast. Key differences include SAGA, the spliceosome and SWI/SNF complexes.

Unlike Drosophila and human, the yeast interactome exhibits a remarkable enrichment of the SAGA complex. Ten of the yeast proteins that we have identified to interact with EWS::FLI1 by Co-IP-MS, are subunits of the SAGA complex. This represents nearly 20% of the identified yeast interactome (n=54) and half of the known *S. cerevisiae* SAGA components (n=20). Moreover, 6 additional SAGA subunits show significant enrichment using a less-stringent cut off (*p*-value < 0.05). In contrast, no SAGA protein was identified to interact with EWS::FLI1 in Drosophila (Molnar *et al*., 2022) and only one, TRRAP (Transformation/transcription domain-associated protein), which can form part of both SAGA and TP60 complexes, has been described to interact with EWS::FLI1 in human cells (Selvanathan *et al*., 2015). Aberrant transcriptional activity by EWS::FLI1 often relies on histone acetyltransferase (HAT) co-activators to remodel chromatin and enhance gene expression. In human cells, the well-characterized EWS::FLI1 interactor p300/CBP serves this function. We hypothesize that the lack of a direct p300 homolog in yeast may facilitate EWS::FLI1 interaction with alternative HAT complexes like SAGA. Although there is currently no direct, substantiated evidence for SAGA’s role as an EWS::FLI1 co-activator in EwS, the strong interaction observed between the oncogene and yeast SAGA subunits raises the possibility of a similar interaction occurring in human cells.

A second notable difference is the absence of spliceosome components in the yeast interactome. The yeast spliceosome contains 84 proteins (Chen & Moore, 2015), 70 of which have orthologues in Drosophila and humans. While many of these orthologues are significantly enriched in the published EWS::FLI1 interactomes of these two animal species (Elzi *et al*., 2014; Molnar *et al*., 2022; Selvanathan *et al*., 2015), none of the 84 yeast spliceosome proteins were detected in the yeast EWS::FLI1 interactome. This lack of interaction in yeast could be due to either the non-conservation of EWS::FLI1 interacting domains in yeast spliceosome proteins or the lower abundance of the spliceosome, which reflects the scarcity of introns and alternative splicing in this model system.

A third striking difference is the absence of SWI/SNF and RSC components in the yeast interactome. The oncogenic effects of EWS::FLI1 are largely mediated through its interaction with the BAF ATP-dependent chromatin remodeling complex. Despite remarkable structural and functional homology between yeast SWI/SNF and human BAF, they present substantial differences both in subunit composition and subunit primary structure (Khavari *et al*., 1993; Muchardt & Yaniv, 1993; Wang *et al*., 1998; Wang *et al*., 1996a; Wang *et al*., 1996b). This is particularly relevant in the case of ARID1A (AT-rich interactive domain-containing protein 1A). ARID1A has been identified as the key BAF complex component mediating the interaction with EWS::FLI1 and acting as a central nucleator of aberrant condensates in EwS cells, specifically through its two N-terminal PrLDs (Prion-Like Domains) which drive dynamic, reversible liquid-liquid phase separation crucial for normal function (Kim *et al*, 2024; Xue *et al*, 2025). Swi1, ARID1A’s yeast homolog, also contains a PrLD, but unlike ARID1A’s PrLDs, Swi1’s PrLD forms stable, self-propagating prion aggregates with a clear “infectious” nature. This distinction may at least in part explain the absence of Swi1, and other SWI/SNF proteins in our yeast EWS::FLI1 interactome.

Our finding that EWS::FLI1 expression in yeast dysregulates only a small number of transcripts is in stark contrast to the hundreds of dysregulated transcripts reported in human cells as well as in Drosophila, which, like yeast, lacks GGAA microsatellites (Hancock & Lessnick, 2008; Kauer *et al*., 2009; Mahnoor *et al*., 2024; Molnar *et al*., 2022). However, a direct comparison between these species is complicated by the different time scales of the experiments. The data from human and Drosophila cells derive from experiments in which the effect of the oncogene took place for extended periods (i.e., days) while our yeast transcriptomes were obtained only 3 and 6 hours after EWS::FLI1 expression was induced. The timing in yeast is well justified for two reasons. Firstly, the 6 hour time point represents about 2-3 consecutive cell cycles which should suffice for EWS::FLI1 to exert its transcription dysregulation effects. Indeed, EWS::FLI1-induced GFP protein expression from the GGAAμSat>GFP transgene, which requires transcription and translation of the GFP mRNA, is readily apparent in 3 hours. Secondly, we wished to avoid unspecific transcriptome changes triggered by the toxic effects of the oncogene which is reflected by the sudden reduction in growth rate that occurs about 9 hours after induction of EWS::FLI1. It is certainly possible that a longer expression time (e.g. 20 hours) may affect transcription of many more genes, but if so, most of the affected genes are likely to be dysregulated by indirect mechanisms and a significant number of them may be a consequence of the toxic conditions brought about by EWS::FLI1 rather than direct targets of the oncogene. Indeed, the 6h EWS::FLI1 yeast signature already contains a substantial number of transcripts encoding for genes known to be expressed in response to different types of cellular stress suggesting that genes dysregulated at this time point may reflect the yeast cells response to EWS::FLI1’s toxic effects.

An alternative and tantalizing hypothesis to explain the small effect of EWS::FLI1 expression on the yeast transcriptome is the lack of conservation of the molecular pathways required for EWS::FLI1 to drive transcriptional changes that are not dependent upon GGAAμSat. Obvious candidates are ETS-TFs, which are unique to animal cells. Outcompetition of ETS-TFs from their biding sites and the resulting dysregulation of the corresponding ETS-TF targets is thought to be one of the main GGAAμSat-independent mechanisms through which EWS::FLI1 reshapes the transcriptional landscape. The yeast genome contains hundreds of potential ETS-TFs binding motifs, but does not encode any ETS-TF protein. Therefore, EWS::FLI1 binding to of potential ETS-TFs binding motifs cannot be expected to exert the profound consequences in transcriptional dysregulation that it has in animal species that contain ETS-TFs. Similarly, the low level of primary sequence conservation or the total lack of other protein complexes linked to EWS::FLI1 (e.g. PcG proteins, NuRD, SWI/SNF, ISWI) may also account for the relatively small effect of the EWS::FLI1 in the yeast transcriptome. This may at least partially explain the lack of correlation between the widespread relocalisation of RNA Pol II to putative ETS-TFs binding motifs, and the relatively low level of transcriptional dysregulation caused by EWS::FLI1. The preferential binding of the 8WG16 antibody to non-phosphorylated RNA Pol II, which may lead to an overrepresentation of non-productive, polymerase binding sites (Miguel *et al*, 2013, 2017), may also contribute to this effect.

To determine whether EWS::FLI1 can relieve transcriptional silencing caused by proximity to GGAAµSats in yeast we have used a rather unconventional approach: instead of making use of minimal promoters as in previous studies (Gangwal *et al*., 2008; Molnar *et al*., 2022) we have used the constitutive promoter CYC1P modified to carry a GGAA repeat insertion upstream. Through this approach we were able, first, to demonstrate that proximity to a GGAA repeat switches down promoter activity, which is consistent with the position effect of the heterochromatic state of the GGAA microsatellite and second, to show that EWS::FLI1 can relieve transcriptional silencing caused by proximity to the GGAA repeat.

The discovery that human EWS::FLI1 can orchestrate gene expression from GGAAμSats in *S. cerevisiae* is remarkable. It is also intriguing since the yeast EWS::FLI1 interactome does not appear to be enriched with SWI/SNF components, and other proteins required for this process in animal cells present substantial evolutionary differences. One of these proteins is CBP/p300. *S. cerevisiae* lacks a direct CBP/p300 protein sequence homolog, but it does have two HATs (histone acetyltransferases), Rtt109 and Gcn5, that share similar structure and function (Li & Shogren-Knaak, 2009; Tang *et al*, 2008). Notably, Gcn5 is part of the yeast SAGA and ADA HAT complexes. Much like CBP/p300, Gcn5 is crucial for activating transcription by acetylating histones, thereby opening the chromatin and making the DNA accessible to the transcription machinery. Indeed, Gcn5 is one of the components of the SAGA complex that is notably enriched in our yeast EWS::FLI1 interactome. Altogether, these observations suggest that in the absence of orthologues to the proteins it interacts with in human cells, EWS::FLI1 is able to mobilise functionally related yeast pathways to drive conversion of closed GGAAμSat chromatin into neoenhancers. If so, it may be possible that the corresponding orthologues, like SAGA and others, may also be utilised by the oncogene in human cells.

## Acknowledgements

We wish to acknowledge the help and support provided by C. Stephan-Otto Attolini and S. Garcia (IRB-Barcelona Bioinformatics and Biostatistics Unit); J.I. Pons, D. Fernández, and F. De Oliveira Monteiro (IRB-Barcelona Functional Genomics Unit); A. Odena, M. Gay, and M. Vilaseca (IRB-Barcelona Mass Spectrometry and Proteomics facility). We thank the Scientific and Technological Centers (CCiTUB), Universitat de Barcelona, and J. Comas for their support and advice on flow cytometry technique. We would also like to thank F. Posas, D. Canadell and M. Nadal (IRB-Barcelona Cell Signalling Group); and J. Ariño and M. Albacar (IBB-UAB) and to all members of our laboratories for very helpful discussions. We are also grateful to the Band of Parents at Hospital Sant Joan de Déu for supporting the overall research activities of the developmental tumor laboratory, PCCB. Work in the González lab is supported by the European Regional Development Fund (ERDF), the Spanish Ministerio de Ciencia e Innovación, the Agencia Estatal de Investigación (grant PID2021- 124716OB-I00), and the Fundació Sant Joan de Déu per la Recerca, Barcelona, Spain.

---

**Supplementary Figure 1.** Growth and expression analysis of the 20×(GGAA)_pCYC1_eGFP reporter strain carrying empty pYES2 or expressing EWS::FLI1, EWS::FLI1_RRLL_, or EWS::FLI1_YS37_.

**(A)** Expression was induced on galactose-containing plates (+GAL). Cells were serially diluted (10-fold) and spotted onto Ura^−^Raff plates without galactose (-GAL, OFF) or supplemented with 2% galactose (+GAL, ON). Growth was monitored for 2 days at 30 °C.

**(B)** The same strains were grown in Ura^−^Raff liquid medium to OD₆₀₀ = 0.2, induced with 2% galactose for 6 h, and harvested. Representative confocal images are shown (left panels): bright-field (top) and GFP fluorescence (bottom). Scale bar, 10 μm.

**(C)** 20×(GGAA)_pCYC1_eGFP strains transformed as in (A) were grown in YNB lacking uracil supplemented with 2% raffinose (Ura^−^Raff) to OD₆₀₀ = 0.25, followed by induction with 2% galactose. Samples were collected prior to galactose addition and after 6 h of induction. Protein extracts were analyzed by Western blot using anti-EWSR1 and anti-GFP monoclonal antibodies to detect EWS::FLI1 variants and GFP, respectively. Arrowheads indicate EWS::FLI1 variants. Ponceau staining is shown as a loading and transfer.

**Supplemental Figure 2.** Known motif analysis of EWS::FLI1 binding sites. Four significantly enriched non-ETS motifs are shown.

**Supplemental Figure 3.** Strategy for genomic insertion of 20x(GGAA) with the modified *pCYC1* promoter regulating eGFP expression. The same approach was used for strains carrying 40x(GGAA) or a random sequence, or by skipping this step in the case of DVS49 (*pCYC1-eGFP::TRP1*). Image generated with SnapGene software.

**Supplemental Figure 4.** Schematic representation of EWS::FLI1 variants cloned into the pYES2 plasmid under *pGAL1* promoter and with the presence of *CYC1* terminator.

## MATERIALS AND METHODS

### Plasmids

The yeast Integrative plasmid (YIp) pRS404, containing the *TRP1* selectable marker (Chee & Haase, 2012) was used to generate the strain DVS050 (*pCYC1*_20x(GGAA)_eGFP) and its derivatives DVS048, DVS049, DVS051(10x(GGAA)) and DVS054(40x(GGAA)). The episomal plasmid pYES2 (Invitrogen) was used for galactose-inducible expression in *S. cerevisiae* under the control of the *GAL1* promoter and carries a *URA3* selectable marker. The pYES2-EWS::FLI1 construct was generated by replacing the *EcoRI-XhoI* fragment in pYES2 with a 1441 bp fragment derived from pUASt-EWS-FLI_1FS_ from (Molnar *et al*., 2022).

To obtain the EWS::FLI1 DNA-binding mutant plasmid (pYES2-EWS::FLI1_RRLL_), pUC-GW-KAN-EWS::FLI1 was linearized by PCR using the mutagenic primers RRLL-Fw and RRLL-Rev. The linearized vector was re-circularized by In-Fusion cloning (Takara Bio), and the *EcoRI-XhoI* fragment containing the mutated open reading frame (ORF) was subcloned into pYES2. All plasmid used in this work are listed in Table S3.

To generate the pYES2-EWS::FLI1_YS37_ plasmid, the *EcoRI-ApaI* fragment of pYES2-EWS::FLI1 was replaced with an 885-nt synthetic DNA fragment (Genewiz) encoding the 37 tyrosine-to-serine substitutions within the EWSR1 prion-like domain.

PCR amplification was performed using KOD DNA polymerase (Merck). In-Fusion cloning (Takara Bio) was carried out using the Snap Assembly kit (Takara). Oligonucleotide primers were synthesized by Sigma-Aldrich, and custom DNA fragments were synthesized by Integrated DNA Technologies (IDT). The sequences of all primers used in this study are listed in Table S4.

### Yeast strains and transformation

The yeast *S. cerevisiae* strain W303-1A (MATa *leu2-3/112 ura3-1 trp1-1 his3-11/15 ade2-1 can1-100*) (Wallis *et al*, 1989) was used to generate the derivatives strains DVS050 (20x(GGAA)), DVS048 (Random_seq), DVS049 (*pCYC1*), DVS051 (10x(GGAA)), and DVS054 (40x(GGAA)) as follows:

To create the DVS050 strain, a 152 bp fragment containing 20x(GGAA) was amplified from the p20x(GGAA)μSat-YFP plasmid (Molnar *et al*., 2022) using the 20x(GGAA)_Fwd and 20x(GGAA)_Rev primers and cloned into the *EcoRI* and *BamHI* sites of pRS426_CYC1p_GFP (Zekhnini *et al*., 2023) to generate pRS426 20x(GGAA)_CYC1p_GFP. A 1399 bp fragment containing 20x(GGAA)_CYC1p_GFP was then inserted into the *BamHI* and *KpnI* sites of the integrative plasmid pRS404. The resulting plasmid was linearized by *Eco81I* and transformed into W303-1A wild-type cells at the *TRP1* locus using the PEG/LiAc transformation method. Transformants were selected on synthetic media lacking tryptophan, and positive clones were verified by PCR.

To obtain DVS048, a random 80-nucleotide sequence (5’-CCAACGCAAAACCTTCACTAGGTTC-TTGACAGTGGAAAAGTGGCCCCGGATTCCTACCGGGTTATCTGAGGATCTGATAA-3’) was synthesized by Integrated DNA technologies (IDT) and inserted into the *BamHI* and *EcoRI* sites of pRS426_CYC1p_GFP. A 1339 bp fragment containing Random_seq_CYC1p_GFP was cloned into the *BamHI* and *KpnI* sites of pRS404, linearized with *Eco81I*, and transformed as described above.

For DVS049, a 1273 bp CYC1p_GFP fragment from pRS426_CYC1p_GFP was cloned into the *BamHI* and *KpnI* sites of pRS404, linearized, and integrated into the genome following the same protocol. All strain used and created in this work are shown in Table S5.

#### Growth of *Escherichia coli* and yeast strains

Unless otherwise indicated, yeast cells were cultured at 28 °C in YP medium (1% yeast extract, 2% peptone) or in synthetic complete (SC) medium lacking the appropriate selection requirements (Adams, 1998). Carbon sources were added as indicated: glucose (Glu; in YPD), raffinose (Raff; in YP-Raff or Ura^-^Raff), or galactose (Gal; in Ura^-^Gal, Ura^-^Raff-Gal, or YP-Gal) at a final concentration of 2%(w/v) unless otherwise stated. Plates contained 2% agar. *E. coli* DH5α cells were used for plasmid propagation and cultured at 37 °C in LB medium, supplemented with 50 μg/ml ampicillin when required. Transformations of *S. cerevisiae* and *E. coli*, as well as standard recombinant DNA techniques, were performed as previously described (Velazquez *et al*, 2020).

### Growth assays

Cells grown on Ura^-^Raff plates were resuspended in sterile water to an optical density at 600 nm (OD_600_) of 0.6 using a Spekol 211 spectrophotometer (Carl Zeiss). Serial 10-fold dilutions of cell suspensions were prepared and spotted onto SC media lacking uracil, complemented with 2% raffinose as a carbon source. When indicated, plates were additionally supplemented with 2% galactose. Growth was monitored for 48-72 hours of incubation.

### Interactome

#### Immunoprecipitation

Immunoprecipitation was performed as previously described, with modifications (Gerace & Moazed, 2014). *S. cerevisiae* DVS050 strain whole-cell extracts expressing EWS::FLI1 for 6 hours were prepared from 100 mL of log-phase culture by bead beating in lysis/IP buffer (50 mM HEPES, pH 7.5; 200 mM sodium acetate; 1 mM EDTA, pH 8.0; 1 mM EGTA, pH 8.0; 0.25% NP40; 1 mM DTT; and protease inhibitor cocktail). The same procedure applied to cells containing empty plasmid was used as a negative control. Three independent biological replicates were performed in each case. Extracts were incubated with anti-EWSR1 antibody (A300-417A, Bethyl Laboratories) crosslinked to Protein A Dynabeads (Thermo Fisher Scientific) using BS3 crosslinker, following the manufacturer’s protocol. The antibody-bead complex was incubated with the lysates for 2 hours at room temperature (RT). Beads were then washed three times with lysis buffer, and bound proteins were eluted by incubating beads with LDS sample buffer. Eluates were resolved by SDS-PAGE (Bolt NuPAGE system, Thermo Fisher) for 5 min, after which the gel band was excised and submitted to the IRB Barcelona Mass Spectrometry & Proteomics Core Facility for mass spectrometry analysis.

#### In-gel Digestion

SDS-PAGE gels lanes were manually excised and cut into cubes. Proteins were reduced and alkylated, and subjected to in-gel-digestion, followed by peptide extraction, as previously described (Shevchenko *et al*, 1996). Peptides extracts were dried using a vacuum concentrator and resuspended in 1% formic acid for subsequent analysis by nanoLC-MS/MS.

#### Reverse Phase Liquid Chromatography Mass Spectrometry

Mass spectrometry was performed on an Orbitrap Eclipse Tribrid mass spectrometer (ThermoFisher Scientific, San Jose, CA) connected to a EVOSEP One (EVOSEP, Odense, Denmark) via nanoEasy Spray Source interface with a stainless steel emitter (EV-1086 EVOSEP). Tryptic peptides (10% of each sample) were loaded onto the EVOTIP (EV-2013 EVOSEP) following the manufacturer’s instructions. The analytical column was a 15cm x 150μm ID, with Dr Maisch C18 AQ, 1.5 μm beads (EV-1137 EVOSEP) with a column oven set at 40°C. The eluents were 0.1% formic acid in water and 0.1% formic acid in acetonitrile. The Evosep One method was 15 SPD (88 minute gradient) (Bache *et al*, 2018), and the flow rate 0.22 μl/min. The mass spectrometer was operated in data independent mode. Full MS1 scans were acquired in the Orbitrap with a scan range of 350 - 1200 m/z and a resolution of 120,000 (at 200 m/z). Automatic gain control (AGC) was set to a target of 1 × 106 and a maximum injection time of 56 ms. MS2 spectra were acquired in data independent mode using 33 m/z variable windows with 0.5 m/z overlap. MS2 spectra were analyzed in the Orbitrap with a resolution of 30,000 (at 200 m/z). A higher energy collision induced dissociation (HCD) method (28% NCE) was applied with an AGC target of 5×104 and a maximum injection time of 55 ms. Orbitrap Eclipse Tune Application v.4.2.4321 and Xcalibur version v.4.7.69.37 were used to operate the instrument and to acquire data.

#### Data Analysis

The spectrum files obtained in the mass spectrometry analyses were searched against a database containing the reviewed Uniprot database for *S cerevisiae* (retrieved in January 2025, 6727 entries), where we added the sequence for EWS::FLI1, using the software Spectronaut v19.6.250122.62635. Acetylation in protein N-terminal, oxidation in Methionine and Methionine excision at protein N-terminal were set as variable modifications, and carbamidomethylation in Cysteine as fixed modification. The maximum number of tryptic missed cleavages and the maximum number of variable modifications allowed were set to 2 and 1, respectively. We used the Proteotypicity Filter with Only Protein Group Specific and the Deep learning-based spectra, RT for the precursor ion generation.

All proteins identified by LC-MS/MS in the negative control (DVS050 strain transformed with the empty plasmid) were removed from the list of proteins identified in the experimental condition (DVS050 strain expressing EWS::FLI1).

The resulting search output was preprocessed, visualized and statistically analyzed using the R 4.3 programming language (https://www.R-project.org/). Protein groups quantified with less than two peptides were filtered out. To ensure robustness, we also required that each protein group had no missing values in at least one experimental condition. We computed protein group abundances from Spectronaut precursor-level abundances using the R implementation (https://github.com/vdemichev/diann-rpackage) of the MaxLFQ (Cox *et al*, 2014) algorithm. Protein group abundances were log2-transformed and the remaining missing values were imputed with normally distributed random numbers with μimputed = μ - 1.8 σ and σimputed = 0.3 σ, where μ and σ are the non-missing values mean and standard deviation, respectively. Data was normalised using cyclic LOESS from limma. The subsequent differential protein abundance analysis was performed using the limma package (Ritchie *et al*, 2015). The linear model was fitted with log₂-transformed MaxLFQ imputed and normalized protein group abundances as the response variable and Condition as the explanatory variable. We tested the contrast FLE vs IgG to identify proteins differentially abundant between these two conditions. Multiple testing correction was applied using the Benjamini & Hochberg method. Finally, we applied standard cutoffs for the fold change (|FC| > 1.5), the adjusted p-value (padj < 0.05) to define significant protein groups.

#### Functional annotation clustering analysis

Functional annotation clustering was performed using the Database for Annotation, Visualization and Integrated Discovery (DAVID 2021). GO_BP_DIRECT terms with a *p-value* of <0.001 were accepted as a significantly enriched.

#### Protein–protein interaction network analysis

Protein–protein interaction (PPI) networks were generated using the STRING v12.0 database (https://string-db.org). The analysis was restricted to the physical subnetwork. Network edges were defined according to confidence scores. Clustering was performed using the *k*-means algorithm. The resulting networks and clusters were visualized using the STRING interface.

#### Identification of yeast orthologs

Human orthologs of yeast EWS::FLI1 interactors were identified using the DRSC Integrative Ortholog Prediction Tool (DIOPT v9.0), using ‘moderate’ or ‘high’ confidence calls, and best score in both forward and reverse searches.

### Transcriptomics

#### RNA-seq sample preparation and sequencing

*S. cerevisiae* DVS050 cells transformed with pYES2, pYES2-EWS::FLI1, or pYES2-EWS::FLI1_RRLL_ plasmids were grown on synthetic media (SC) lacking uracil with 2% raffinose to OD_600_ = 0.15. Then, galactose was added to 2% final concentration. Cells were harvest after 3 or 6 hours by centrifugation (3,000 x*g*, 5 minutes). Total RNA extraction was done by hot phenol (Latorre *et al*, 2022). The quality of the extracted total RNA was assessed using a 2100 Bioanalyzer system (Agilent, 5067- 1511) with an RNA integrity score (RIN) > 8. The concentration of total RNA extractions was quantified with the Nanodrop One (Thermo Fisher), and RNA integrity was assessed with the RNA ScreenTape assay of the Tapestation 4200 platform (Agilent). Libraries for RNA-Seq were prepared at IRB Barcelona Functional Genomics Core Facility. Briefly, mRNA was isolated from 1.3 μg of total RNA and used to generate dual-indexed cDNA libraries with the Illumina stranded mRNA ligation kit (Illumina) and UD Indexes Set A (Illumina). Ten cycles of PCR amplification were applied to all libraries.

Libraries were quantified using the Qubit dsDNA HS assay (Invitrogen) and quality controlled with the Tapestation HS D5000 assay (Agilent). An equimolar pool was prepared with the eighteen libraries and submitted for paired-end 150 nt sequencing on a NovaSeq6000 S4 (Illumina). 102 Gbp of reads were produced, with a minimum of 14.6 millions of paired-end reads per sample.

#### RNA-seq data processing

Adapters from reads were removed using Trim Galore v.0.6.7 (https://github.com/FelixKrueger/TrimGalore) with default parameters. Resulting reads were aligned to the *S. cerevisiae* R64 genome using STAR v.2.7.10a (Dobin *et al*, 2013). The count matrix was generated in R (https://www.R-project.org) through the function “featureCounts” of the Rsubread package (Liao *et al*, 2014).

#### Differential expression analysis

Differential expression was performed using the DESeq2 package (Love *et al*, 2014). For plotting purposes and generating principal components, the count matrix was normalized using the “vst” function. Genes with log2FC-shrinked > |1| and pval.adj<0.05 were considered differentially expressed. Three biological replicates of each experimental condition were analysed.

#### Gene Ontology and KEGG pathway analysis

Functional annotation of Gene Ontology (GO) Biological Process terms and KEGG pathways was performed using the online tool Database and Annotation, Visualisation and Integrated Discovery (DAVID 2021). GOTERM_BP_3 and KEGG_PATHWAY terms with a *p-value* of <0.05 were accepted as a significantly enriched.

### ChIP-seq

#### Sample preparation

ChIP-Seq was performed as previously described (de Jonge *et al*, 2020) with minor modifications. DVS050 cells carrying either empty pYES2 plasmid (control) or pYES2-EWS::FLI1 were cultured in Ura^-^ Raff medium to an to OD_600_ = 0.15, after which galactose was added to a final concentration of 2% (w/v). After 6 hours, cells were harvested and crosslinked by addition of 37% formaldehyde to final concentration of 2%, followed by incubation for 6 min at 25°C with gentle shaking (70 rpm). The crosslinking reaction was quenched by adding 4.5 M Tris-HCl (pH 8.0) to a final concentration of 1.5 M, followed by incubation for 2 min at 25°C with gentle shaking (70 rpm). Cells lysis was performed using glass beads (G8772, Sigma-Aldrich), and chromatin was fragmented using a Bioruptor Pico sonicator (Diagenode) for 9 cycles (30 s on / 30 s off), yielding DNA fragments of 200-500 bp. Chromatin extracts were incubated overnight at 4°C with 4 μg of either anti-EWSR1 antibody (sc-48404, Santa Cruz Biotechnology), anti-RNA polymerase II antibody (sc-56767, Santa Cruz Biotechnology), or anti-GFP antibody (11814460001, Roche) as an IgG control. The following day, 25 μl of Dynabeads Protein G (Thermo Fisher Scientific) were added to the antibody-chromatin mixtures and incubated for 2h at RT. Beads were collected using magnetic rack and washed twice with 1 ml of lysis buffer containing 350 mM NaCl, once with wash buffer (10 mM Tris-HCl (pH 8.0), 250 mM LiCl, 0.5% NP-40, 0.5% sodium deoxycholate, 1 mM EDTA) and once with 1 ml TE buffer (10 mM Tris-HCl, 1 mM EDTA, pH 8.0). Beads were resuspended in 94 μl TE/SDS buffer (10 mM Tris-HCl, pH 8.0; 1 mM EDTA; 1% SDS) and incubated overnight at 65°C. The next day, 2 μl of RNAse A/T1 (Thermo Fisher) were added and incubated for 30 min, followed by addition of 20 μl of Proteinase K (20mg/ml) and incubation for 2h at 37 °C. DNA was purified using the QIAquick MinElute PCR Purification Kit (Qiagen), according to the manufacturer’s instructions. All chromatin immunoprecipitation experiments were performed in three independent biological replicates.

#### Sequencing

The concentration of the DNA samples (inputs and IPs) was quantified with Qubit dsDNA HS kit and fragment size distribution was assessed on a TapeStation 4200 using the D5000 HS assay (Agilent). Libraries for ChIP-Seq were prepared at IRB Barcelona Functional Genomics Core Facility. Briefly, dual-indexed DNA libraries were generated from 1 ng of DNA samples using the NEBNext Ultra II DNA Library Prep kit for Illumina (New England Biolabs). 15 cycles of PCR amplification were applied to all libraries. The final libraries were quantified using the Qubit dsDNA HS assay (Invitrogen) and quality controlled with the Tapestation D5000 HS assay. An equimolar pool was prepared with the twenty-four libraries for Single End 50 nts sequencing on a NextSeq2000 (Illumina). More than 23 Gbp were produced, with a minimum of 15.5 million of single-end reads per sample

#### Data pre-processing and quality control

Paired-end FastQ files adapters were removed using Rfastp. Reads were aligned to the *S. cerevisiae* W303 genome with Bowtie2 v2.5.4 using --local -N 1 -k 1. Quality control of FastQ and BAM files was performed with FastQC v0.12.1 and MultiQC. BigWig coverage tracks for IGV were generated with deepTools bamCoverage with effectiveGenomeSize = 12,100,000 and BPM normalization (bins per million mapped reads).

#### Peak calling

Peaks were calling against inputs using MACS2. Peaks were annotated with HOMER v2 using Saccharomyces_cerevisiae.R64-1-1.60_v2.gtf genome annotation.

#### Consensus peaks

Consensus peaks were called with DiffBind package v3.16 and defined as those present in at least one sample of a condition to retain condition-exclusive peaks. For each consensus peak, the presence in each sample was annotated considering the threshold presence/absence by signal>0.1. Visualization of peaks was performed with DeepTools (i) scale-regions (peaks stretched to gene regions) and (ii) reference-point (genomic position of peaks) with 3000 upstream and 3000 downstream of the reference point.

#### Motif enrichment analysis

Peak BED files for each condition were used as targets, and consensus peaks were used as background. HOMER v2 function FindMotifsGenome.pl was performed with default size = 200 bp.

#### Genome-wide motif search

Genome-wide searches for GGAA repeats were performed in *S. cerevisiae* using R v4.2.1 (https://www.R-project.org) with the Bioconductor packages BSgenome and Biostrings, loading the UCSC sacCer3 genome assembly. Exact matches (forward and reverse strand) were identified with the matchPattern() function.

### Microscopy and flow-cytometry

Cells were grown on synthetic media (SC) lacking uracil with 2% raffinose (Ura^-^Raff) up to OD_600_ ≈ 0.2. Then, galactose was added to reach a concentration of 2%, after 6 hours samples were subjected to confocal microscopic observation and images were acquired with a SP8 Leica confocal image microscope and processed in ImageJ.

Flow cytometry assays were carried out following the same procedure but using 96-well plates. Samples were analyzed by Cytek® Aurora (4-laser and 64 Fluorescence Emission Detection Channels). The full spectrum (365–829 nm) of 10.000 yeast cells were recorded from each sample according to their FSC and SSC distributions and unmixed to identify the GFP fluorescence signal. DVS049 cells expressing GFP were used as positive control and W303-1A wild-type cells were used to set zero GFP signal. Cytometry data were analyzed using FlowJoTM Software (BD Life Sciences) and expressed as mean percentage of GFP fluorescence or histograms of GFP signal intensity.

### Immunoblotting

Cells were grown in Ura^-^Raff to early exponential phase (OD_600_ ≈ 0.2), then 2% galactose was added. Cultures were normalized to an OD_600_ = 0.8; 10 ml of cells were harvested by centrifugation and frozen at -80 °C. Thawed samples were treated in 0.1 M NaOH for 15 minutes on ice, pelleted, resuspended in 100 μL of 2x LDS sample buffer, and boiled 10 min. Lysates were cleared by centrifugation at 10,000 x*g* and 10 μL were loaded in Bis-Tris SDS-PAGE (Bolt® 4-12% gradient; Invitrogen) and transferred to a nitrocellulose membrane. Membranes were blocked in 1.5% powdered milk in PBS-T during 1h 30min at RT, then incubated over-night at 4 °C primary antibodies: anti-EWSR1 (Santa Cruz Biotechnology, 1:500), and anti-GFP (Invitrogen, 1:500). Primary antibody was removed and membranes washed 1×15 minutes and 1×5 minutes in PBS-T and incubated with Alexa Fluor®790 or Alexa Fluor®680 secondary antibodies (1:2500, Jackson ImmunoResearch). Membranes were washed 1×15 minutes, 1×10 minutes and 3×5 minutes in PBS-T. Last wash step was performed with PBS buffer. Results were quantified using the Odyssey® M Imaging System (Li-Cor) and LI-COR imaging software.

### Statistical analyses

Statistical analyses were performed using GraphPad Prism 9.2 for macOS X (GraphPad Software, La Jolla, CA, USA; www.graphpad.com). p-values were calculated using one-way ANOVA for GFP fluorescence levels and two-way ANOVA for western blot band intensities.

