## Supplemental Figures 1, 2, 3, 4 and 5 for "Ancient and Animal-Specific Regulatory Modes of EWS::FLI1 Revealed by a Minimal Yeast Model"

Suppl. Figure 1

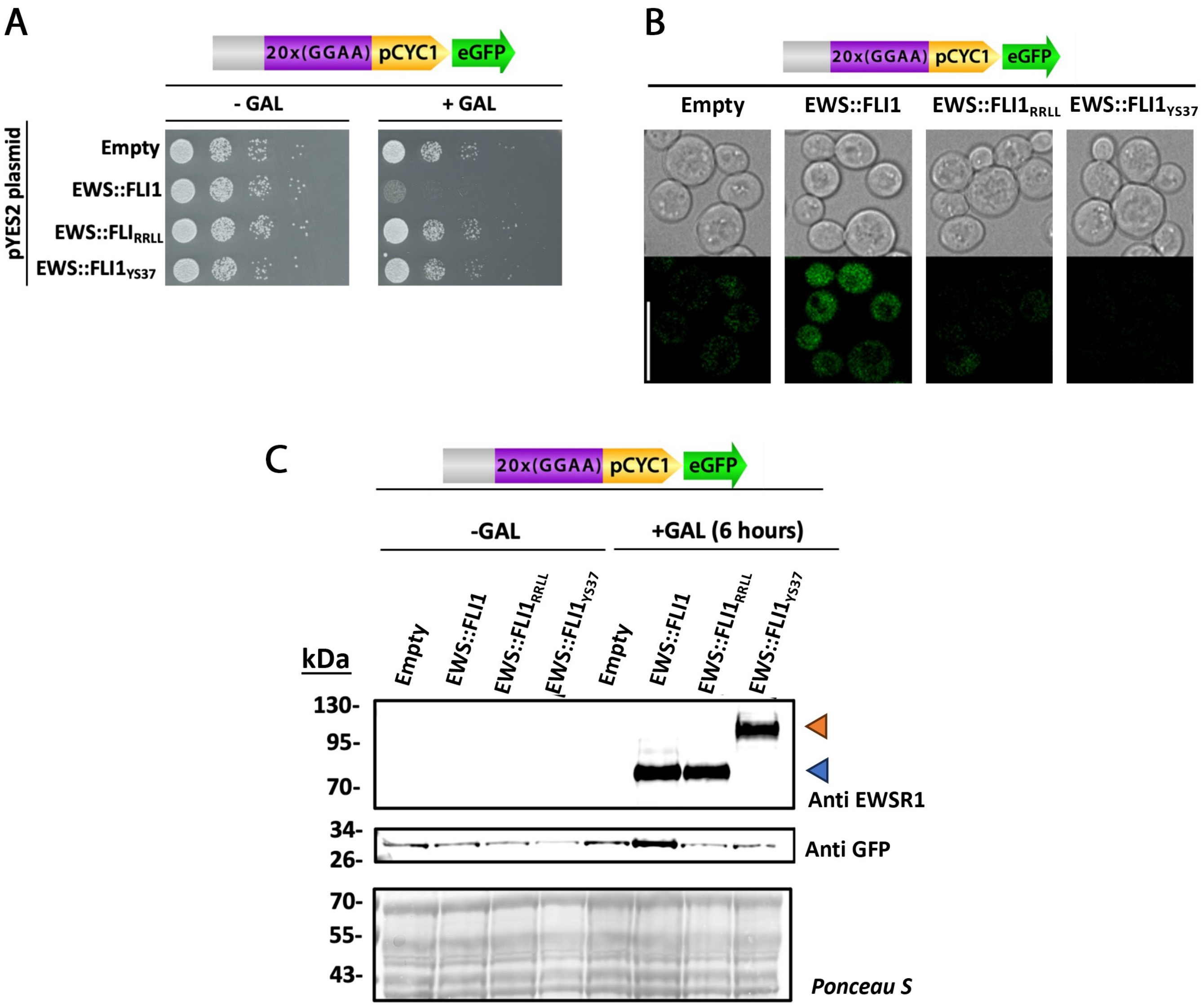

Suppl. Figure 2

| EWS::FLI1 known motif (non-ETS motifs) | P-value | % Targets | % Background | Name |
|---|---|---|---|---|
| 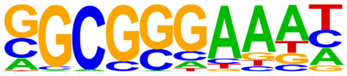 | 1e-49 | 27.52% | 9.64% | E2F4 |
| 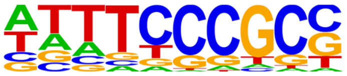 | 1e-37 | 19.79% | 6.45% | TFDP1 |
| 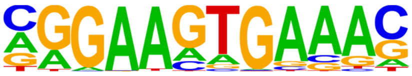 | 1e-35 | 49.88% | 29.32% | PU.1-IRF |
| 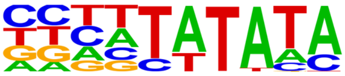 | 1e-23 | 41.10% | 25.08% | TATA-box |

Suppl. Figure 3

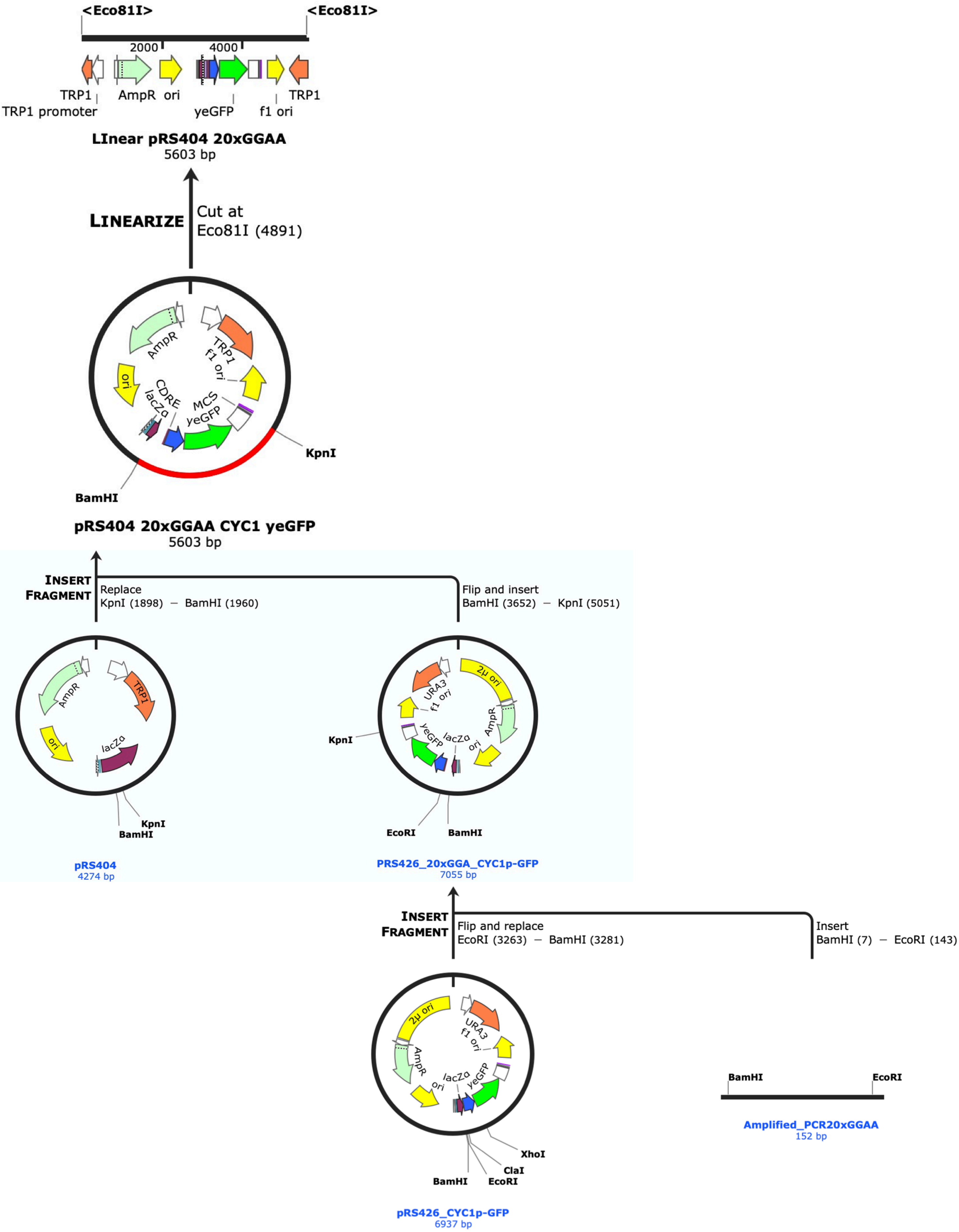

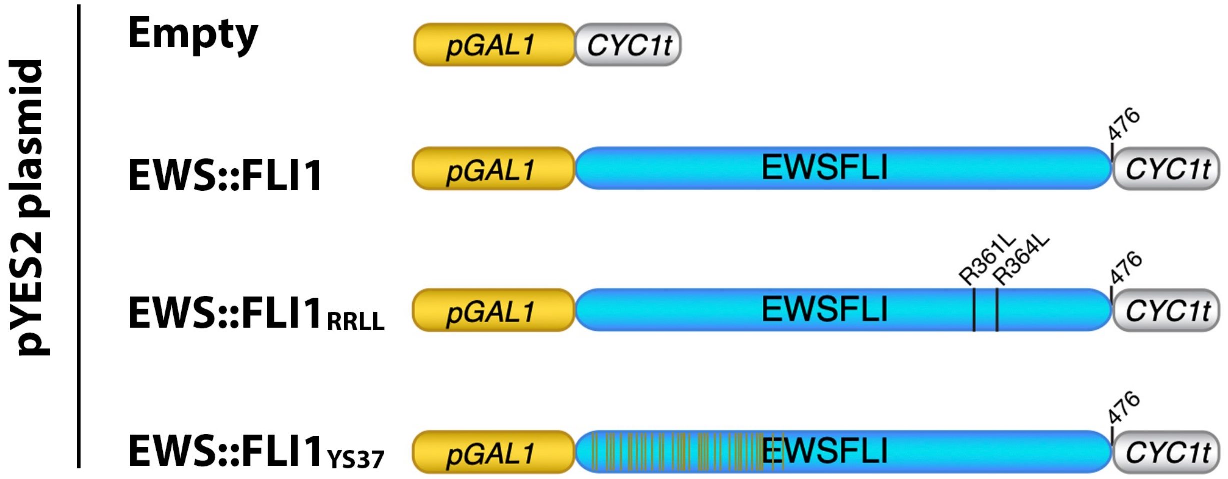
pYES2 plasmid
Empty
EWS::FLI1
EWS::FLI1RRLL
EWS::FLI1YS37
pGAL1
CYC1t
EWSFLI
476
R361L
R364L
