## Supplementary material for "Ancient and Animal-Specific Regulatory Modes of EWS::FLI1 Revealed by a Minimal Yeast Model": Table S3, S4 and S5

### Table S3. Plasmids used in this study

| Plasmid | Description | Source/Reference |
|---|---|---|
| **pRS426_CYC1p-GFP** | Centromeric plasmid to express GFP under modified *CYC1* promoter | (Zekhnini *et al.*, 2023) |
| **pRS426_10x(GGAA)_CYC1p-GFP** | Centromeric plasmid to express GFP with a 10x(GGAA) upstream | This study |
| **pRS426_20x(GGAA)_CYC1p-GFP** | Centromeric plasmid to express GFP with a 20x(GGAA) upstream | This study |
| **pRS426_40x(GGAA)_CYC1p-GFP** | Centromeric plasmid to express GFP with a 40x(GGAA) upstream | This study |
| **pRS426_RandomSeq_CYC1p-GFP** | Centromeric plasmid to express GFP with a RandomSeq upstream | This study |
| **pRS404** | Yeast integrative plasmid (YIp) *TRP1* marker | (Chee & Haase, 2012) |
| **pRS404_10x(GGAA)_CYC1p-GFP** | YIp plasmid with a 10x(GGAA)_CYC1p driving GFP expression | This study |
| **pRS404_20x(GGAA)_CYC1p-GFP** | YIp plasmid with a 20x(GGAA)_CYC1p driving GFP expression | This study |
| **pRS404_40x(GGAA)_CYC1p-GFP** | YIp plasmid with a 40x(GGAA)_CYC1p driving GFP expression | This study |
| **pRS404_RandomSeq_CYC1p-GFP** | YIp plasmid with a RandomSeq_CYC1p driving GFP expression | This study |
| **pYES2** | Episomal plasmid for galactose inducible expression | Invitrogen |
| **pYES2 EWS::FLI1** | EWS::FLI1 wild-type expression | This study |
| **pYES2 EWS::FLI1$_{RRLL}$** | EWS::FLI1 DNA binding mutant expression (R361L+R364L) | This study |
| **pYES2 EWS::FLI1$_{YS37}$** | EWS::FLI1 with 37 Tyr residues mutated to Ser in the EWSR1 portion | This study |

### Table S4. Oligonucleotides used in this study

| Oligo name | Sequence | Use |
|---|---|---|
| **20xGGAA Fw** | GCTAGCGGATCCAAGCTT | Integrate 10, 20 and 40x(GGAA) in genome |
| **20xGGAA Rev** | GCTAGAATTCCGCTCGCTAGAGTCTCCG | Integrate 10, 20 and 40x(GGAA) in genome |
| **yeGFP comp Rev** | CCGTAAGTAGCATCACCTT | To check integrations in genome |
| **pYES EWS-FLI Fw** | GCTCGGATCCACTAGTAAC | EWS::FLI1 cloning |
| **pYES EWS-FLI Rev** | AGGGACCTAGACTTCAGG | EWS::FLI1 cloning |
| **RRLL-Fw** | CTCGCCCTCCTATACTATTACGACAAGAAC | EWS::FLI1 R361L + R364L mutagenesis |
| **RRLL-Rev** | GTATAGGAGGGCGAGGGACAGCTTGTC | EWS::FLI1 R361L + R364L mutagenesis |

**Table S5. *Saccharomyces cerevisiae* strains used in this study**

| Name | Genotype | Source / Reference |
|---|---|---|
| **W303-1A** | MATa *leu2-3/112 ura3-1 trp1-1 his3-11/15 ade2-1 can1-100* | (Wallis *et al.*, 1989) |
| **DVS048** | W303-1A Randomseq_*pCYC1_eGFP::TRP1* | This study |
| **DVS049** | W303-1A *pCYC1_eGFP::TRP1* | This study |
| **DVS050** | W303-1A 20x(GGAA)_*pCYC1_eGFP::TRP1* | This study |
| **DVS051** | W303-1A 10x(GGAA)_*pCYC1_eGFP::TRP1* | This study |
| **DVS054** | W303-1A 40x(GGAA)_*pCYC1_eGFP::TRP1* | This study |
